# Probabilistic data integration identifies reliable gametocyte-specific proteins and transcripts in malaria parasites

**DOI:** 10.1101/199356

**Authors:** Lisette Meerstein-Kessel, Robin van der Lee, Will Stone, Kjerstin Lanke, David A Baker, Pietro Alano, Francesco Silvestrini, Chris J Janse, Shahid M Khan, Marga van de Vegte-Bolmer, Wouter Graumans, Rianne Siebelink-Stoter, Taco WA Kooij, Matthias Marti, Chris Drakeley, Joseph J. Campo, Teunis JP van Dam, Robert Sauerwein, Teun Bousema, Martijn A Huynen

## Abstract

*Plasmodium* gametocytes are the sexual forms of the malaria parasite essential for transmission to mosquitoes. To better understand how gametocytes differ from asexual blood-stage parasites, we performed a systematic analysis of available ‘omics data for *P. falciparum* and other *Plasmodium* species. 18 transcriptomic and proteomic data sets were evaluated for the presence of curated “gold standards” of 41 gametocyte-specific versus 46 non-gametocyte genes and integrated using Bayesian probabilities, resulting in gametocyte-specificity scores for all *P. falciparum* genes.

To illustrate the utility of the gametocyte score, we explored newly predicted gametocyte-specific genes as potential biomarkers of gametocyte carriage and exposure. We analyzed the humoral immune response in field samples against 30 novel gametocyte-specific antigens and found five antigens to be differentially recognized by gametocyte carriers as compared to malaria-infected individuals without detectable gametocytes. We also validated the gametocyte-specificity of 15 identified gametocyte transcripts on culture material and samples from naturally infected individuals, resulting in eight transcripts that were >1000-fold higher expressed in gametocytes compared to asexual parasites and whose transcript abundance allowed gametocyte detection in naturally infected individuals. Our integrated genome-wide gametocyte-specificity scores provide a comprehensive resource to identify targets and monitor *P. falciparum* gametocytemia.

## Introduction

Despite a decrease in malaria incidence and mortality over the past two decades, malaria remains a major global health challenge ^1,2^. Furthermore, the emergence and spread of insecticide resistance in mosquitoes ^3^ and artemisinin resistance in *Plasmodium falciparum* (*Pf*) ^4–6^ threaten recent gains in malaria control. The decline in malaria burden and the necessity to contain artemisinin-resistance have increased interest in malaria elimination that may require interventions that specifically aim to prevent malaria transmission. Malaria transmission depends on male and female gametocytes, the sexually reproducing forms of the *Plasmodium* parasite that are ingested by blood-feeding *Anopheles* mosquitoes. In the mosquito gut, gametocytes may complete the parasite’s reproductive cycle and, following sporogonic development, render the mosquito infectious. Factors that govern gametocyte production and infectivity remain poorly understood. Whilst recent studies have shed light on the processes controlling gametocyte commitment ^7,8^, commitment and maturation of gametocytes may differ between infections and over the course of infections, under influence of environmental and host factors ^9,10^. A better understanding of gametocyte dynamics during infections, as well as the development of tools to monitor or target gametocytes, may be informed by high-throughput protein and transcriptome studies ^11,12^. In the past 15 years, a number of large-scale studies on *Plasmodium* gametocytes have been reported: the proteome of *Pf* and the rodent malaria parasite *Plasmodium berghei (Pb)* have been examined by mass spectrometry ^13–21^, and the transcriptome of both species by microarray and RNA sequencing ^15,20,22–28^. These studies differed in their focus and resolution in examining (sexual) developmental stages and each faced challenges in detecting low abundance proteins ^29^ and by the purity of parasite populations ^13,14^. The use of fluorescent parasites and fluorescence-assisted sorting of staged parasites have recently permitted a better discrimination of proteins in either male or female gametocytes ^16,17,20^ and have allowed more detailed comparisons of *Plasmodium* life-stages. However, individual studies are still vulnerable to imperfect sample purity, and other sources of uncertainty such as correct gene identification for accurate peptide assignment. These technical and methodological challenges lead to discrepancies between individual studies and hamper firm conclusions about gametocyte-specificity of proteins and transcripts.

We utilized the numerous published proteomics and transcriptomics *Plasmodium* data sets in a comprehensive data integration framework to obtain a consensus of gametocyte-specific transcripts and proteins. Our data integration approach is an adaptation of the naïve Bayesian classifiers that have previously been applied in the prediction of protein interactions and components of cellular systems ^30,31^. The framework calculates probabilities that any given transcript or protein is gametocyte-specific given the evidence presented across the total of transcriptomics and proteomics data. A key aspect of the methodology is that it takes into account the predictive power of each contributing data set: it assigns weights to data sets based on their ability to distinguish gold standard lists of gametocyte and asexual proteins. These we have constructed using existing literature where life-stage specificity was confirmed using classical “non-omics” approaches (e.g. protein detection in immunofluorescence-assays, functional/genetic studies), followed by expert curation. The most informative data (from datasets with the highest discriminative power against the gold-standard lists) will thus contribute most to the predictions, while less informative data are down-weighted. This allows for (i) the resolution of conflicting evidence without disregarding data, and (ii) the construction of a transparent scoring system in which the relative contribution of each data set is directly visible.

Using this approach, we propose a robust gametocyte-specificity score for all *Pf* genes that allows a consensus list of gametocyte-specific genes at protein and transcript level. We illustrate the utility of our findings by examining naturally acquired responses to newly identified gametocyte-specific proteins in gametocyte-carriers and non-carriers by protein microarray. In addition, we confirmed gametocyte-specificity for a selection of gametocyte-specific transcripts using culture material from geographically distinct *Pf* strains and samples from naturally infected malaria patients.

## Results

### Weighted integration of proteomics and gene expression data

Using Bayesian statistics, we integrated *Plasmodium* mass spectrometry and transcript datasets from 18 different studies (Table 1) on *P. falciparum (Pf;* n=14), *P. berghei (Pb;* n=3) and *P. vivax (Pv*; n=1). Since gametocyte biology differs between *Plasmodium* species, scores were calculated for the total set of *Plasmodium* studies and for *Pf* only. Unsupervised clustering of genes based on peptide counts or mRNA expression resulted in grouping according to data acquisition method rather than parasite stage (Fig 1), illustrating the necessity of a supervised approach to discriminate between gametocyte-specific and non-gametocyte genes. To objectively assess the value of individual data sets and allow their assembly into a gametocyte-specificity score, we created a gold standard that served as a benchmark for every sample. This gold standard was collected from literature review and comprises two lists; one of asexually expressed proteins, mainly blood stage but also sporozoite and liver stage, and one of known gametocyte proteins (Supplementary Table S1). A gametocyte-specificity score was then derived for each gene by comparing its expression in all studies to the relative expression of the gametocyte and asexual gold standards in those samples (Supplementary Fig S1-2). Proteins or RNAs detected in a study with high discriminative power for gametocyte and asexual gold standard genes (ratio of gametocyte to asexual gold standard genes) received higher gametocyte-specificity scores than those detected in a study with lower discriminative power. The individual log-transformed scores per gene were combined for proteomics and transcriptomics data separately. Scores for *Pf-*only studies (Supplementary Table S2) and all combined data sets were highly correlated (Fig 2D; Pearson’s r=0.9867 and r=0.9514 for proteomics and transcriptomics, respectively) and the latter were used in the remainder of the manuscript. The distribution of scores for proteins and transcripts are presented for the two sets of genes of the gold standard as well as for all other genes (Fig 2A). As expected, the gametocyte and asexual gold standard set of genes are perfectly separated by their respective proteomics-derived scores and show only little overlap in their transcriptomics-derived scores (Fig 2B). The shift in the density peak of the proteomics compared to transcriptomics is due to an inherent property of the method that gives a negative score for gametocyte-specificity to all proteins that were not detected in the proteomics studies (n=1583). The highest scoring 100 genes for proteomics and transcriptomics contained 26 (63.4 %) and 15 (36.6 %) of the 41 gold standard gametocyte genes, respectively (Fig 2C), indicating both the respective discriminating power of the gold standard and that many other genes are as specific for gametocytes as the highest gold standard representatives. Translationally repressed genes are common in late stage female gametocytes ^32,33^ and are detectable by high transcriptomics and low proteomics score. In our analysis 461 genes have this profile (Supplementary Table S3), including genes that are known to be translationally repressed like Pfs28 ^32,34^ and 186 genes with a previously reported bias towards expression in female gametocytes ^20^.

**Figure 1.**
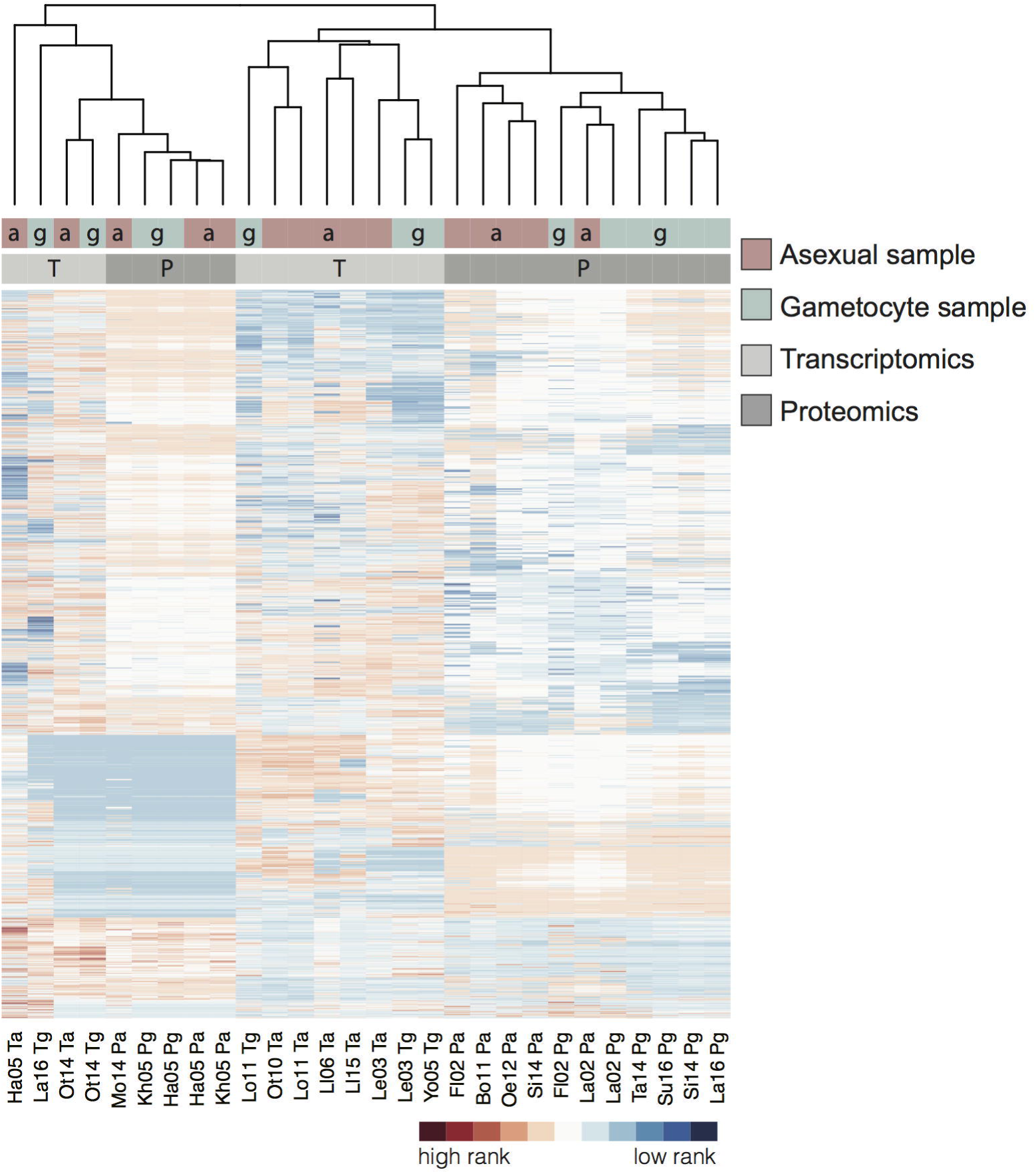
Clustered data sets used in this study with genes ranked according to their protein or transcript expression. Level of expression as detected in the respective samples with unique peptide counts for MS data and percentiles for transcriptomics. The studies are clustered using complete linkage according to their overall gene expression similarities (Euclidean distance). See Table 1 for study keys. Distribution of asexual(a)/gametocyte(g) samples (red/blue) is shown in top bar, proteomics (P) and transcriptomics (T) (dark/light grey) in lower bar.

**Figure 2.**
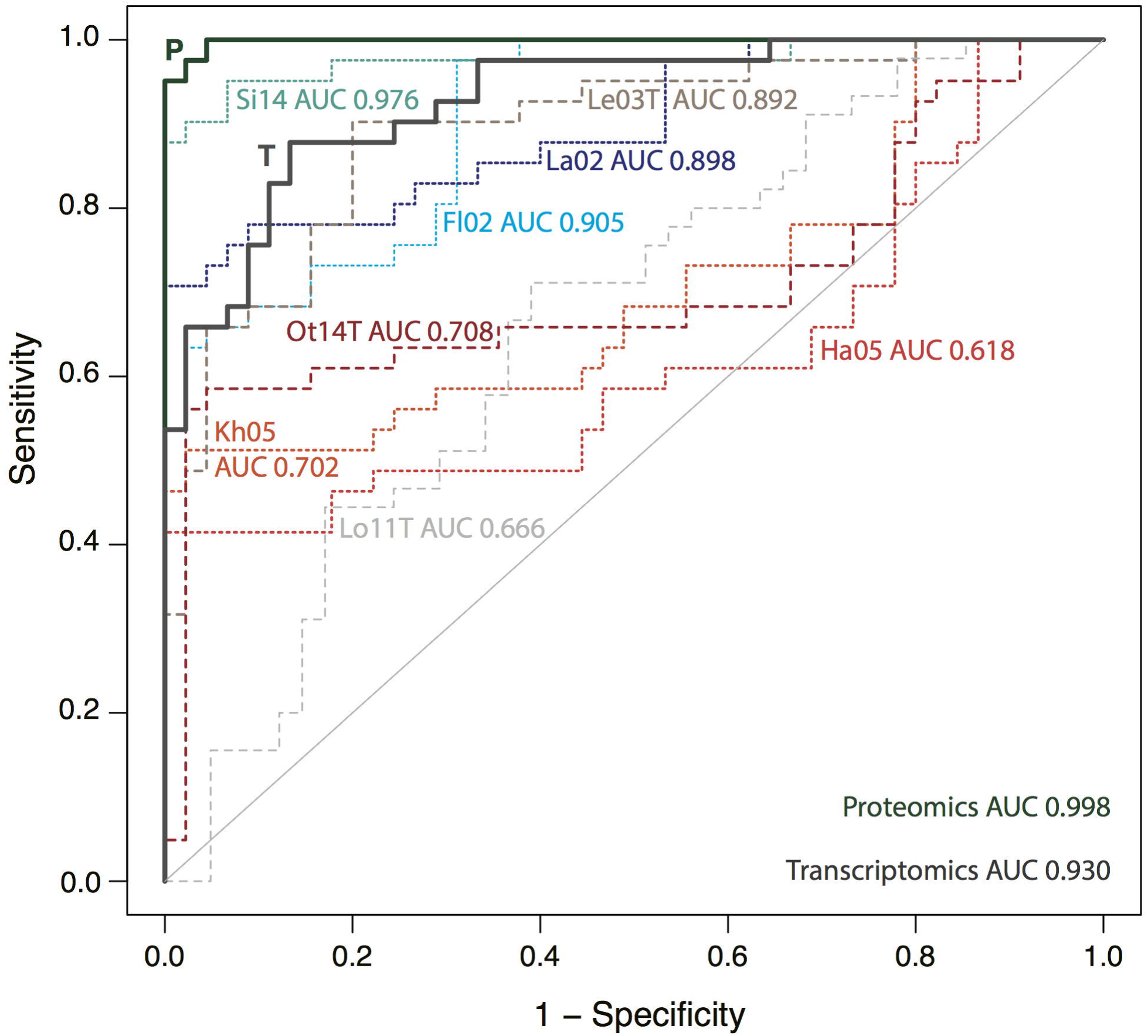
Gametocyte-specificity scores for P. falciparum genes derived from proteomics (P) and transcriptomics (T) data sets. (A) Boxplot for integrated scores for the two gold standard sets and all other Pf genes, derived from proteomics, transcriptomics or all data sets (combined). (B) Density of P and T gametocyte scores, individual gold standard genes and their scores are indicated at the bottom (red, asexual, blue gametocyte). (C) 100 highest ranking proteins and transcripts, gametocyte gold standard in blue. (D) Correlation of the gametocyte-specificity scores derived from all integrated MS studies and *Pf* MS studies only.

**Table 1.**
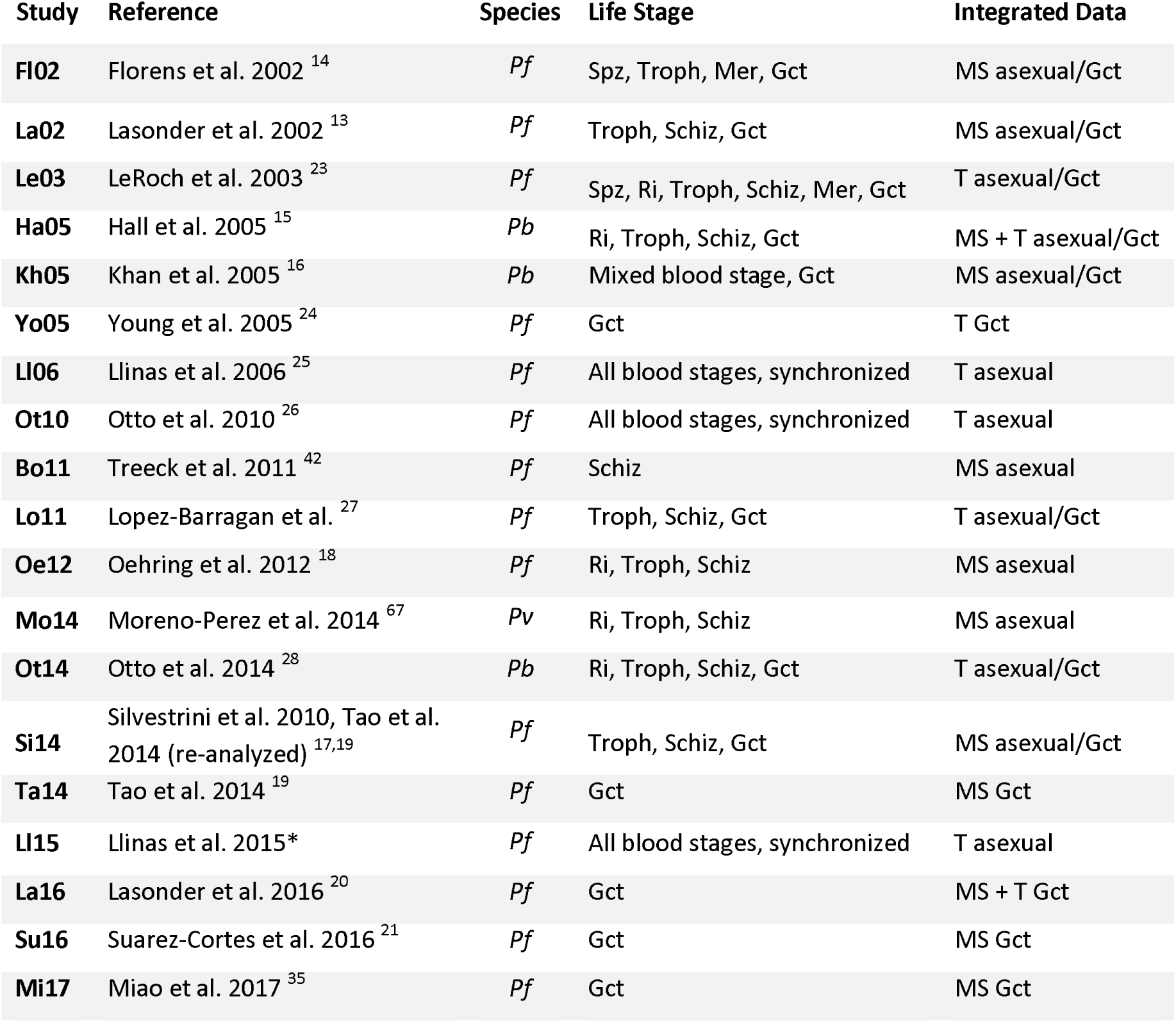
Data sets used for integration. Life stages Spz sporozoites, Ri rings, Troph trohozoites, Schiz schizonts, Mer merozoites, Gct gametocytes. MS mass spectrometry (highest unique peptide count in any of the samples), T transcriptomics (highest percentile in any of the samples). Data from Miao et al. 2017 was integrated after analyses of high scoring proteins, ranks and scores are included in Supplementary Table 2. *Asexual microarray data by Llinas and others retrieved from plasmoDB version 28 (Data set “Pfal3D7 real-time transcription and decay”), no accompanying publication.

| Study | Reference | Species | Life Stage | Integrated Data |
| --- | --- | --- | --- | --- |
| FI02 | Florens et al. 2002 <sup>14</sup> | *Pf* | Spz, Troph, Mer, Gct | MS asexual/Gct |
| La02 | Lasonder et al. 2002 <sup>13</sup> | *Pf* | Troph, Schiz, Gct | MS asexual/Gct |
| Le03 | LeRoch et al. 2003 <sup>23</sup> | *Pf* | Spz, Ri, Troph, Schiz, Mer, Gct | T asexual/Gct |
| Ha05 | Hall et al. 2005 <sup>15</sup> | *Pb* | Ri, Troph, Schiz, Gct | MS + T asexual/Gct |
| Kh05 | Khan et al. 2005 <sup>16</sup> | *Pb* | Mixed blood stage, Gct | MS asexual/Gct |
| Yo05 | Young et al. 2005 <sup>24</sup> | *Pf* | Gct | T Gct |
| LI06 | Llinas et al. 2006 <sup>25</sup> | *Pf* | All blood stages, synchronized | T asexual |
| Ot10 | Otto et al. 2010 <sup>26</sup> | *Pf* | All blood stages, synchronized | T asexual |
| Bo11 | Treeck et al. 2011 <sup>42</sup> | *Pf* | Schiz | MS asexual |
| Lo11 | Lopez-Barragan et al. <sup>27</sup> | *Pf* | Troph, Schiz, Gct | T asexual/Gct |
| Oe12 | Oehring et al. 2012 <sup>18</sup> | *Pf* | Ri, Troph, Schiz | MS asexual |
| Mo14 | Moreno-Perez et al. 2014 <sup>67</sup> | *Pv* | Ri, Troph, Schiz | MS asexual |
| Ot14 | Otto et al. 2014 <sup>28</sup> | *Pb* | Ri, Troph, Schiz, Gct | T asexual/Gct |
| Si14 | Silvestrini et al. 2010, Tao et al. 2014 (re-analyzed) <sup>17,19</sup> | *Pf* | Troph, Schiz, Gct | MS asexual/Gct |
| Ta14 | Tao et al. 2014 <sup>19</sup> | *Pf* | Gct | MS Gct |
| LI15 | Llinas et al. 2015* | *Pf* | All blood stages, synchronized | T asexual |
| La16 | Lasonder et al. 2016 <sup>20</sup> | *Pf* | Gct | MS + T Gct |
| Su16 | Suarez-Cortes et al. 2016 <sup>21</sup> | *Pf* | Gct | MS Gct |
| Mi17 | Miao et al. 2017 <sup>35</sup> | *Pf* | Gct | MS Gct |

### Cross-validation illustrates the improved predictive power of the integrated data

Tenfold cross-validation was performed using random subsamples of the gold standard lists to predict the ranks of left-out genes. The resulting proteomics ranking shows near perfect sensitivity, with all but two gold standard gametocyte genes ranking higher than the gold standard asexual genes (Fig 3). The added value of our integrated approach is illustrated by the receiver operating characteristic curve where the integration of data sets gave higher sensitivity and areas under the curve for both proteomics and transcriptomics than any individual study (Fig 3). Using the Bayesian integration based on the complete gold standards, we ranked all *Pf* proteins by giving them a gametocyte-specificity score (Supplementary Table S2). All proteins with a score >5 (n=602) were considered gametocyte-specific. Most of these have not consistently been described as “specific” or “enriched” in gametocytes in the original data sets (Fig 4A and Supplementary Table S4). Previous studies defined 315 ^13^ to 1725 ^20^ proteins as gametocyte-specific for *Pf*. Not only did our integrated approach lead to a better recovery of gold standard listed known gametocyte proteins, we also identified 178 genes with undescribed function as gametocyte-specific (Supplementary Table S2). We further identify a number of proteins as gametocyte-specific even though they had been reported as asexual by previous studies (Fig 4B). A recent proteomics study of male and female *Pf* gametocytes ^35^, not included in our original analysis, was used to test the robustness of our scores. When we included this data set in our final Bayesian proteomics scores, both gametocyte scores and gene ranks before and after addition of this data set were highly correlated (Pearson’s r=0.997 and Spearman’s rho=0.995, respectively). Furthermore, the top 100 gametocyte proteins did not change and the top 602 proteins were 96% identical (578 of 602). Taken together, cross-validation and independent data suggest that the integrated gametocyte-specificity score is robust and contains potential novel gametocyte markers.

**Figure 3.**
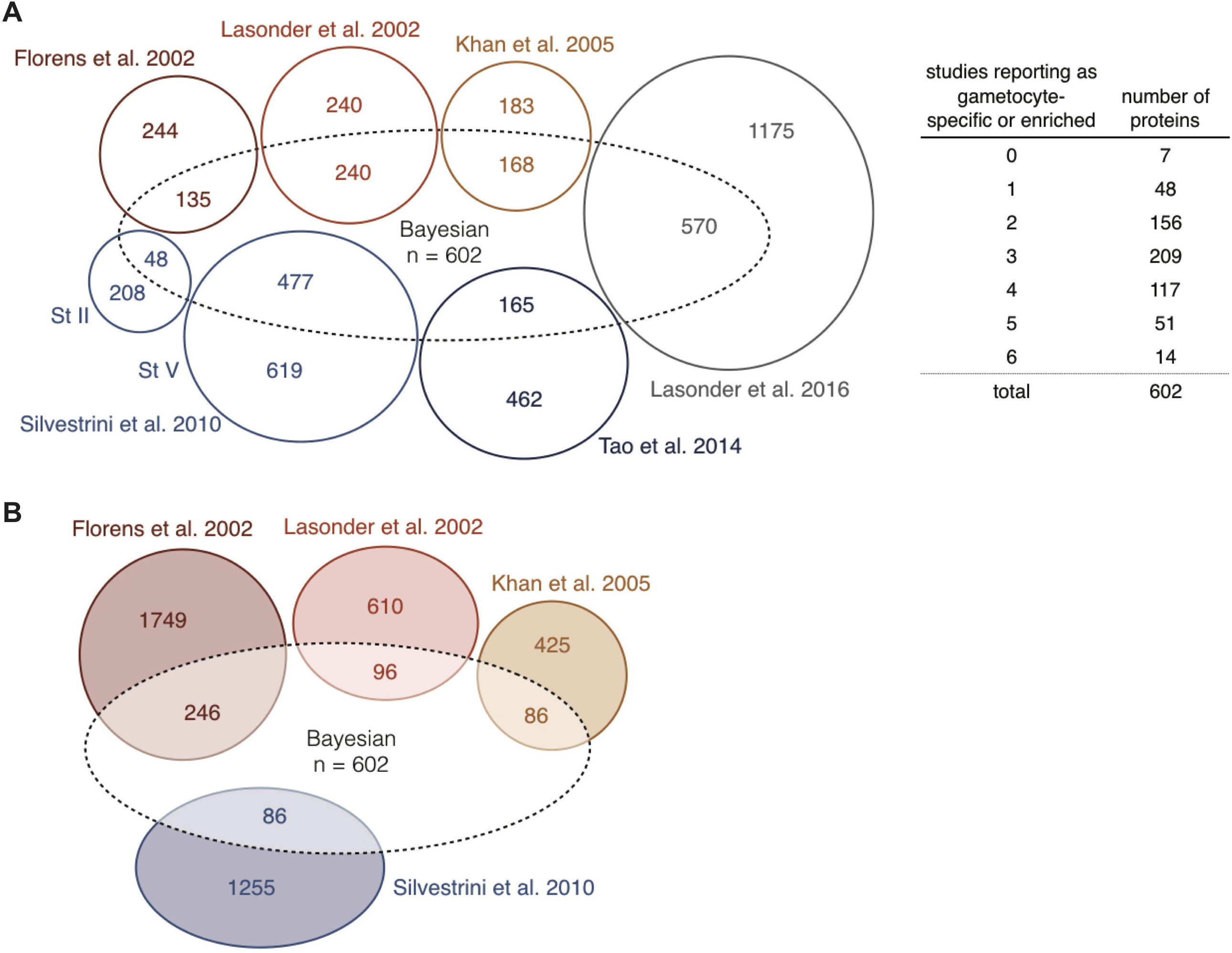
Validation of Bayesian gametocyte scoring with area under the curve (AUC) values. Integrated data and individual data sets are compared by 10-fold cross-validation (subsampling of gametocyte and asexual gold standard sets). Integrated proteomics (P) and transcriptomics (T) scores in bold lines. P. berghei data sets in shades of red, individual proteomics and transcriptomics studies with short and long dashes, respectively. See Table 1 for study keys.

**Figure 4.**
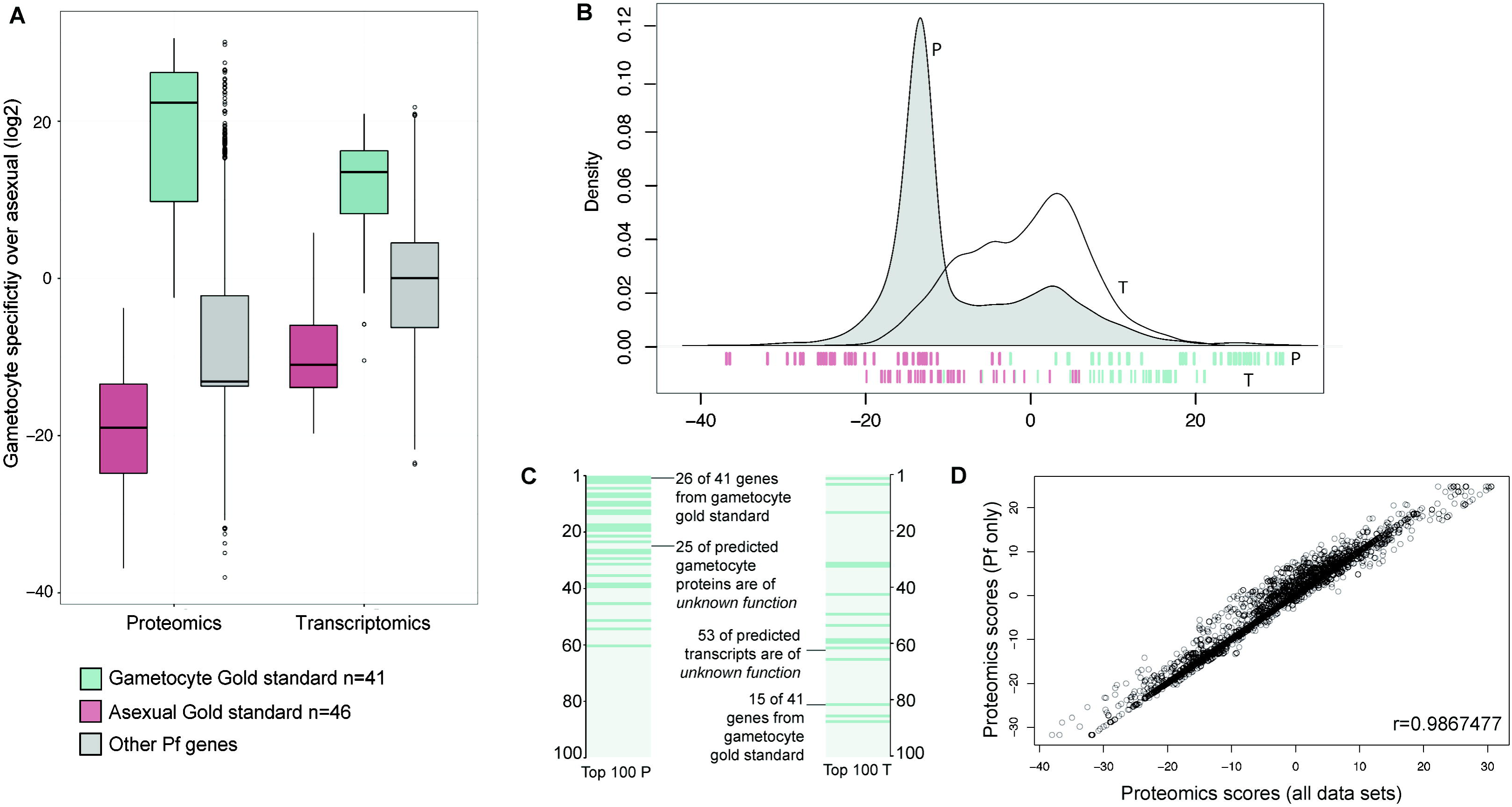
Comparison of reported gametocyte-specific proteins in mass spectrometry studies. (A) Proteins reported as gametocyte-specific by six individual studies, agreements on gametocyte-specificity are summarized in the table. Bayesian: gametocyte-specific proteins (n=602) that have a score > 5 after data integration. The overlap with previously published data sets is shown, but not to scale. Overlap between the individual studies is not shown for better visibility. Note that the Lasonder 2002 study includes proteins that were found in gametocytes or gametocytes and gametes. (B) Proteins that were reported as non-gametocytic and are (partially) included after data integration

### Predicted gametocyte-specific proteins are recognized by gametocyte-carriers

As an illustration of the utility of gametocyte-specific proteins as markers of gametocyte exposure, we utilized protein microarray data from a study that aimed to characterize the immune profile associated with transmission-reducing immunity in naturally infected gametocyte carriers (Stone, Campo *et al.* 2017 accepted manuscript ^36^). For the current study, we compared responses to our gold standard gametocyte genes (n=40) and novel gametocyte genes from our 100 highest scoring proteins that were on the array (n=30). Antibody prevalences for these genes were compared between Gambian gametocyte carriers and Gambians who carried asexual parasites but not gametocytes as determined by microscopy. Antibody responses to the predicted gametocyte-specific proteins were significantly higher in gametocyte carriers (p=0.005), while for the gold standard antigens this difference was less significant (p=0.058) (Fig 5A, Mann-Whitney U test). When antigens were analysed individually, a significantly higher antibody prevalence in gametocyte carriers was detected for five novel gametocyte antigens (Fig 5B, p<0.05 in Fisher’s exact, corrected for multiple testing, Supplementary Table S5, Supplementary Fig S3). Only two of these five have an assigned function – a DNA ligase, and Gamete egress and sporozoite-traversal protein (GEST). For two of the three remaining *Plasmodium* proteins, we were able to predict a function based on homology, using the sensitive homology detection tool HHpred ^37^. PF3D7_1251000 is homologous to the co-chaperone HSP20 heat shock protein and PF3D7_1439600 is homologous to the MLRQ subunit of complex IV of the oxidative phosphorylation, underlining the enrichment in mitochondrial proteins as discussed below (Supplementary Table S6 includes homology predictions for all conserved, highly gametocyte-specific *Plasmodium* proteins).

**Figure 5.**
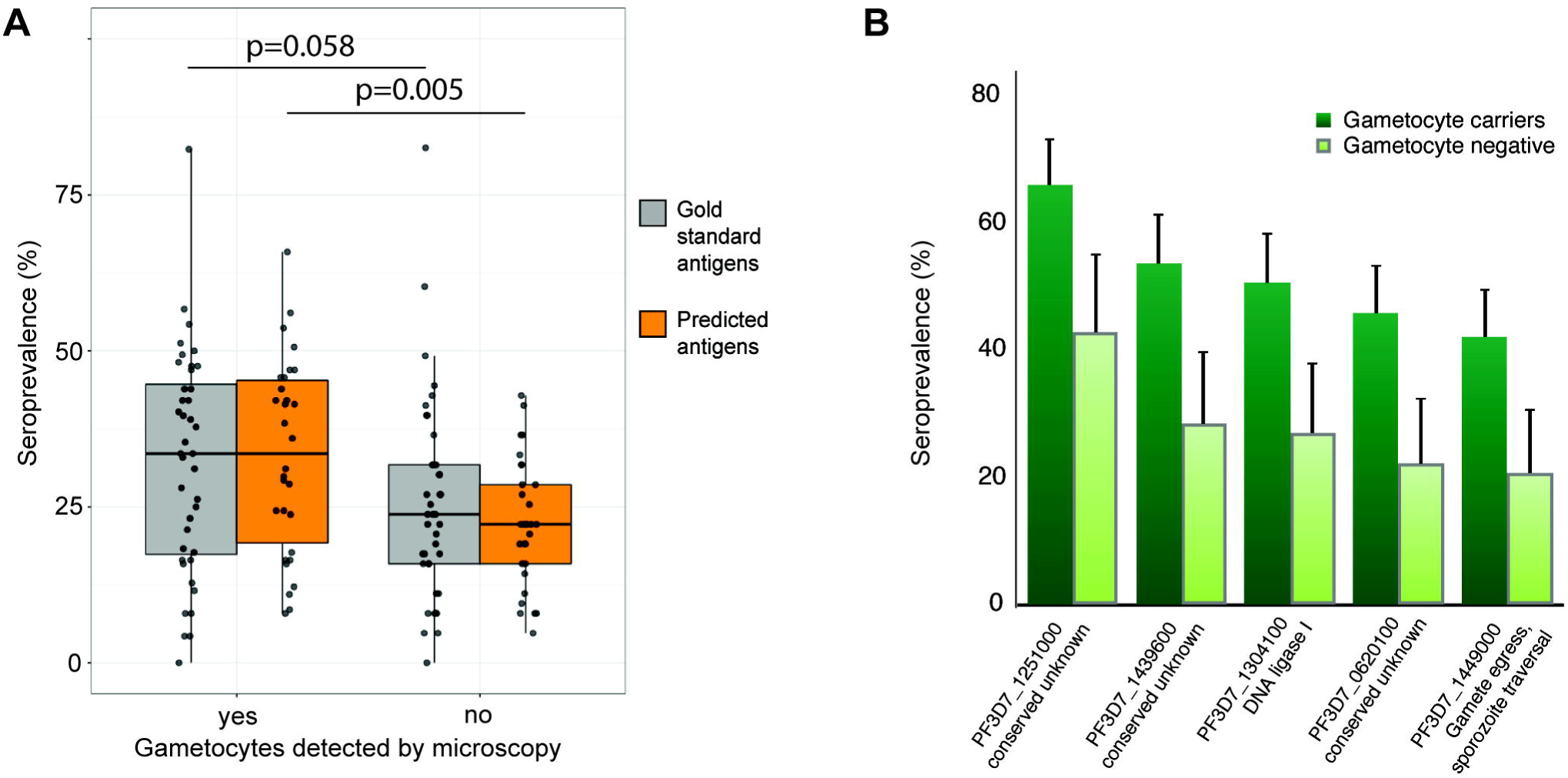
Seroprevalence in two cohorts of parasite carriers in The Gambia. (A+B) Antibodies against the highest scoring gametocyte-specific proteins were measured on protein microarrays. Comparison of positivity (mixture-model cutoff) in gametocyte carriers (n=164) and non-carriers (n=63). Gametocyte presence determined by microscopy. All individuals were positive for asexual parasites. (A) Prevalence of antigens from the gold standard (n=40) and predicted gametocyte-specific proteins (n=30), Mann-Whitney U test (B) Antigens of five predicted gametocyte-specific proteins are preferentially recognized by gametocyte carriers. Error bars indicate the upper limit of the 95% confidence interval around the proportion. p<0.05 Fisher’s exact test, corrected for multiple testing of a total of 70 antigens (Benjamini-Hochberg)

### Gametocyte-specific RNA transcripts detect (submicroscopic) gametocyte carriage

Of the 100 highest-scoring transcripts, 15 non-gold standard candidates were selected for qRT-PCR validation based on their gametocyte scores in a preliminary analysis (Table 2). Mature gametocytes of four *Pf* strains from different geographical origins were compared to asexual blood stage parasites. The minimum transcript abundance difference (Ct_Asexuals_-Ct_Gametocytes_) ranged from 4.76 to 14.95 (Fig 6A and Supplementary Table S7 qPCR & primers), reflecting 27.1 to 31,500-fold higher transcript numbers in gametocytes compared to asexual parasites and confirming pronounced upregulation of all selected targets in gametocytes. With a very conservative threshold of 1,000-fold enrichment in three of the four strains tested (Fig 6A), eight of the 15 tested transcripts were highly specific to gametocytes. Transcript abundance in ring-stage parasites was assessed and compared to Pfs25 mRNA, an established and highly abundant yet intron-less female gametocyte specific transcript ^11,38^. Five out of eight gametocyte specific transcripts were undetectable in asexual ring stages at ≤10^5^ parasites/mL, similar in specificity to Pfs25 (Fig 6B); the five most sensitive gametocyte markers detected gametocytes across the range of 10^2^ – 10^6^ gametocytes/mL (Fig 6C). In RNA samples from a previously reported clinical trial conducted in Kenya ^39^, all eight gametocyte markers detected gametocytes at densities below 10^3^/mL (Fig 6D).

**Figure 6.**
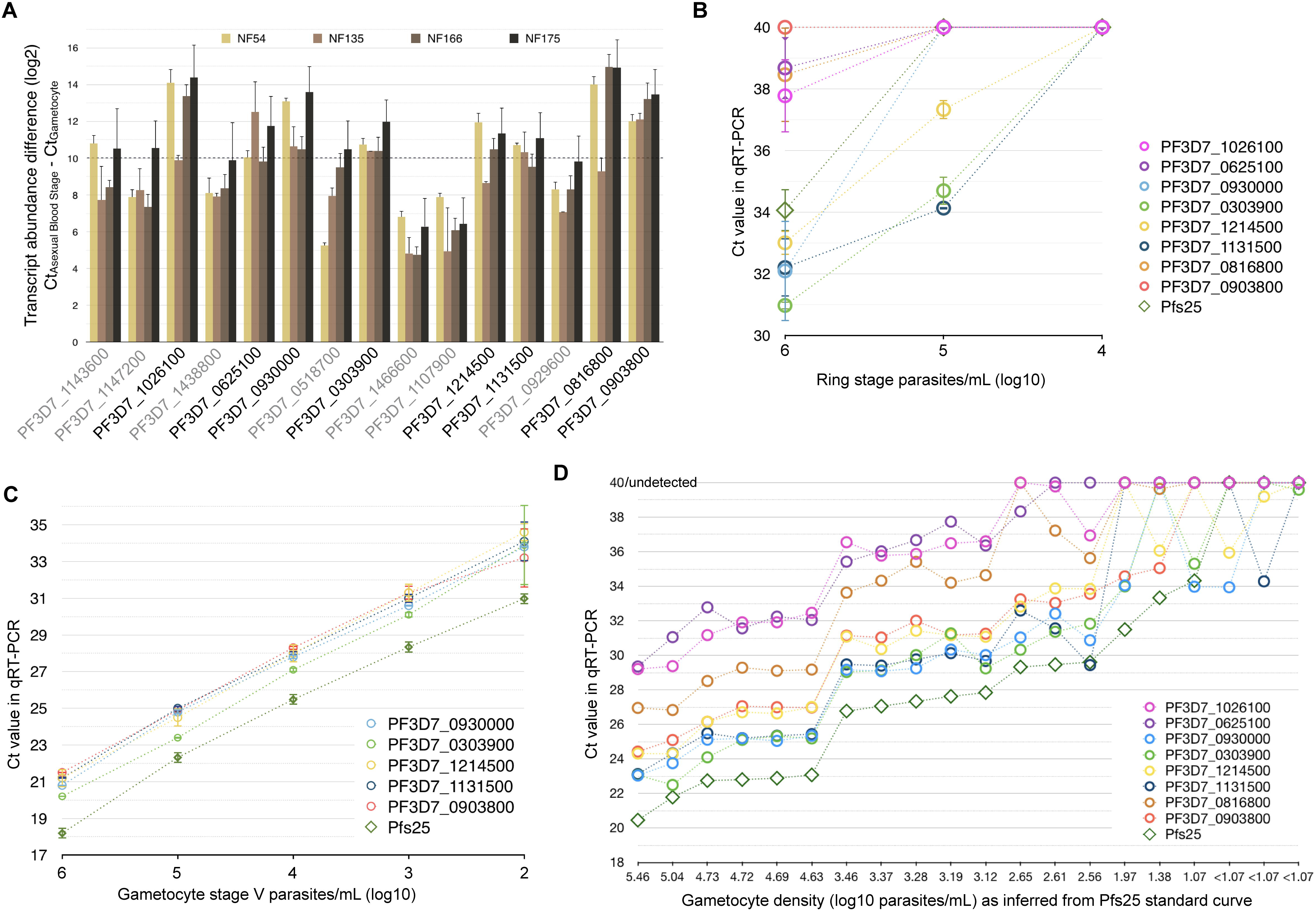
Validation of gametocyte-specific targets in qRT-PCR. Targets are sorted for decreasing gametocyte-specificity in all panels, see Table 2. (A) Minimum transcript abundance in blood stage versus gametocytes in different Pf strains. 1000-fold enrichment of transcript in gametocytes over asexuals was assumed when delta-Ct was 10 or higher (dashed line), considering the lowest Ct value detected in any asexual concentration-matched sample. This threshold was not met by the transcripts with gene IDs in grey. (B) Detection limit of eight validated targets alongside Pfs25 in serial dilutions of Pf NF54 asexual stage parasites (ring stage parasites 10-20 hours post invasion). (C) Detection limit of the most sensitive targets in serial dilutions of stage V gametocytes. (A-C) For Pf NF54, all n=3, other strains n=2 biological replicates (error bars: standard error of the mean), all measurements in triplicates. (D) Sensitivity of eight validated targets in Kenyan blood samples of varying gametocyte densities.

**Table 2.** Properties of putative gametocyte-specific targets. Rank in transcriptomics (all data sets) for specificity in gametocytes. Random sample of top 100, excluding the gold standard. If primers are not intron-spanning, samples were DNase I treated. Ct difference is the difference between the lowest Ct detected in asexual samples and the highest Ct in concentration-matched stage V gametocytes, averaged across strains Pf NF54, NF135, NF166 and NF175.

| Rank | Gene ID | Description | Name | Intron-spanning | Least Ct difference |
| --- | --- | --- | --- | --- | --- |
| 3 | PF3D7_1143600 | conserved Plasmodium protein, unknown function | — | no | 9.37 |
| 5 | PF3D7_1147200 | tubulin--tyrosine ligase, putative | — | no | 8.52 |
| 6 | PF3D7_1026100 | conserved Plasmodium protein, unknown function | — | yes | 12.94 |
| 7 | PF3D7_1438800 | conserved Plasmodium protein, unknown function | — | yes | 8.57 |
| 8 | PF3D7_0625100 | sphingomyelin synthase 2, putative | SMS2 | no | 11.04 |
| 12 | PF3D7_0930000 | procollagen lysine 5-dioxygenase, putative | — | no | 11.96 |
| 14 | PF3D7_0518700 | mRNA-binding protein PUF1 | PUF1 | yes | 8.31 |
| 15 | PF3D7_0303900 | phosphatidylethanolamine-binding protein, putative | — | yes | 10.87 |
| 16 | PF3D7_1466600 | conserved Plasmodium protein, unknown function | — | no | 5.67 |
| 17 | PF3D7_1107900 | mechanosensitive ion channel protein, putative | MSCS | no | 6.33 |
| 18 | PF3D7_1214500 | conserved Plasmodium protein, unknown function | — | yes | 10.61 |
| 24 | PF3D7_1131500 | conserved Plasmodium protein, unknown function | — | no | 10.41 |
| 51 | PF3D7_0929600 | G2 protein, putative | — | yes | 8.37 |
| 61 | PF3D7_0816800 | meiotic recombination protein DMC1, putative | DMC1 | yes | 13.29 |
| 75 | PF3D7_0903800 | LCCL domain-containing protein | CCp4 | yes | 12.70 |

### Gametocyte-specific proteins are enriched for cytoskeletal movement and metabolism functions

To uncover novel characteristics underlying gametocyte function, we analyzed over-represented gene ontology (GO) terms in our integrated consensus gametocyte proteins. The 100 highest ranked proteins were examined for enrichment of GO terms that reflect specific biological functions. Microtubule based movement, metabolism of carboxylic acids and metabolism of nucleic acids were highly enriched among gametocyte proteins (Supplementary Fig S4). Four out of the six putative *Pf* dynein heavy chain proteins are found back among the 100 most gametocyte-specific proteins, alongside a tubulin gamma chain and a tubulin chaperone. The importance of DNA elongation and ligation processes is reflected in GO term associations as well as in antibody response to a DNA ligase (Fig 5B). The “classic” GO term enrichment calculation was complemented by a rank-based gene set enrichment analysis (GSEA). GSEA uses all *Pf* genes and their respective (proteomics derived) gametocyte-specificity score and thus does not include an arbitrary cutoff of the proteins that are or are not gametocyte-specific. It confirmed the above-mentioned results, and in addition to those the terms “mitochondrial protein complex” (GO:0098798) and “TCA” (GO:0006099) were enriched, stressing both mitochondrial location and processes. Although male gametocytes carry pre-synthesized proteins to rapidly form eight motile gametes upon activation in the mosquito midgut, we did not observe flagellum associated terms in the GSEA. The reason for this is that current GO term annotation for *Pf* has only one gene with the “cilium” GO term for cellular component (GO:0005929; PF3D7_1025500), one for “axoneme” (GO:0005930; PF3D7_0828700) and none with the biological-process terms “cilium or flagellum-dependent cell motility” (GO:0001539) or “axoneme assembly” (GO:0035082). We supplemented the GO annotation with a list of 28 *Pf* cilium genes (Methods). The newly assembled “cilium” GO term now acquired the highest enrichment score in the GSEA (Supplementary Fig S5). This may reflect the formation of the flagella of the microgamete but may also (partially) reflect intracellular trafficking and or be associated with genome replication as the term has overlap with the genes annotated for microtubule processes (Supplementary Table S5 GO terms).

## Discussion

Combining proteomic and transcriptomic data from 18 sources, we present an integrated consensus score for gametocyte-specific proteins and transcripts. We predict 602 gametocyte-enriched proteins of which 186 are currently without ascribed function. We illustrate the potential utility of our gametocyte score by providing evidence for differential recognition of gametocyte proteins by naturally infected gametocyte carriers and the sensitive detection of mRNA of novel gametocyte transcripts in field samples.

The gametocyte proteome of *P. falciparum (Pf)* has been assessed repeatedly. Individual lists of gametocyte-specific proteins ^13,14,16,17,19,20^ have unavoidable limitations related to comparator (asexual) parasite stages, sample purity, assay sensitivity and arbitrary cut-offs used to define gametocyte-specificity and show only partial agreement. To acquire a more robust gametocyte-specificity score, we integrated data from these individual studies, along with studies of purified asexual parasites and related *Plasmodium* species. Including gene expression data from multiple species generally increases the likelihood that the combined gene expression data reflect underlying biology, as observed in the *Apicomplexa* ^40,41^. We applied a Bayesian classifier first applied to ‘omics data by Jansen and colleagues ^30^ and adapted by Van der Lee and colleagues to identify genes involved in anti-viral immune responses ^31^. The probabilistic approach combines the evidence from all studies in an unbiased way, without giving *a priori* preference of one study over another. Instead, the measurements of all studies were weighted inherently during the scoring process by assessing the retrieval of a gold standard set of genes. As these gametocyte and asexual gold standard sets are of central importance to the study, they have undergone expert curation (see Methods and Acknowledgments). The power of this integrative approach lies not only in weighting data sets by the retrieval of gold standard genes but also in the opportunity to exclude proteins from the gametocyte-specific list by appreciating their presence in (other) asexual samples. A further strength of the approach is that it allows the ranking of gametocyte proteins that have only been reported in a subset of studies. Our integration of data sets reveals that 602 proteins are likely to be specific to gametocytes although very few gametocyte-specific proteins were detected in every underlying dataset and seven proteins had never before been reported as gametocyte-specific. A general limitation of all mass spectrometry (MS) studies is their bias toward highly abundant proteins. Proteins with low-level expression may be missed in a bulk proteome analysis. After integration of the MS studies listed in Table 1, 1583 *Pf* proteins were never detected, representing approximately 28% of all proteins encoded by *Pf*. Some of these might be of too low abundance or expressed during sporozoite or gamete, ookinete and liver stage, which are underrepresented or not included in our data, respectively. New advances in MS that include the sensitive detection of peptides from currently understudied *Plasmodium* life stages may shed light on these currently uncharacterized genes. In addition, approaches that focus specifically on post translational modifications like phosphorylation of proteins as has been done for asexual parasites ^42–45^ may add new lines of evidence towards gametocyte-specific functions of proteins. Our approach suggests that the currently available MS data is sufficiently comprehensive to identify stage-specific proteins when analysed in an integrative approach. We examined this directly by incorporating a new *Pf* gametocyte MS study ^35^ in our scoring. The authors reported 44 new gametocyte-specific proteins that were not reported by earlier studies. We compared this data set to our integrated data set and found 24 of the 44 had been detected in one or more erythrocytic stages or sporozoites ^14,18,42,46,47^ while 11 others had been identified in a (single) gametocyte sample before (Supplementary Fig S6). Importantly, the scores and top 100 gametocyte genes remained unaltered by integrating this new dataset.

The ranking of gametocyte-specificity that we provide here can i) aid in understanding the biology of this life stage and ii) improve diagnostics related to gametocyte exposure and carriage. Regarding gametocyte biology, our high-ranking gametocyte-specific genes are enriched for mitochondrial, metabolism and microtubule processes and DNA replication, supporting the quality of the data integration. The enrichment of mitochondrial localization and process is consistent with what we know about the enlarged mitochondrion of gametocyte stages ^48^ and increased activity of the citric acid cycle ^49^. DNA replication terms are highly enriched which is consistent with what happens in the subsequent life stage in which the (micro)gamete rapidly duplicates its genomic DNA three times. Regarding the use of the gametocyte score to inform gametocyte diagnostics, diagnostics can directly detect nucleic acids specific to gametocytes ^11^ or detect antibody responses reflecting past/recent exposure as is increasingly used for asexual *P. falciparum* and *P. vivax* parasites ^50,51^. We use our integrated gametocyte list to explore its utility for both approaches. We validated 15 transcript targets in four different *Pf* strains, comparing transcript abundance in gametocytes and asexual parasites. All tested targets were enriched in gametocytes. Five targets were tested for their sensitivity and can recognize 100 gametocytes/mL, while the signal is undetectable when fewer than 10^5^-10^6^ ring-stage parasites/mL are present. In practical terms, these markers may be used to reliably detect gametocytes at densities well below the microscopic threshold of detection in samples without high-densities of asexual parasites, similar to the gametocyte marker that is currently most widely used, the female gametocyte-specific Pfs25 ^38^.

As an alternative approach to the detection of gametocyte carriage in populations, we utilized a gametocyte-enriched protein microarray (Stone, Campo *et al*. 2017 accepted manuscript ^36^) to determine antibody responses to genes that we here describe as highly gametocyte-specific. The bacterial expression system used for the array has known limitations with the expression of conformational proteins ^52^ and should thus be considered a ‘rule in’ rather than ‘rule out’ approach to immune recognition. Moreover, the array was constructed with the aim of detecting surface proteins or exported proteins whilst our list does not require these characteristics. Only 30 of our top 100 novel gametocyte antigens were thus printed on this array. Antibody responses to five gametocyte proteins were significantly more prevalent in gametocyte carriers than in carriers of the asexual blood stage only. This is the first evidence that antibody responses may be indicative of current gametocyte carriage. Importantly, the dichotomization of gametocyte-exposed and non-exposed individuals was based on a single time-point screening for gametocytes by microscopy. Microscopy has a low sensitivity for detecting gametocytes that commonly circulate at low densities ^53^ and several of the asexual parasite carriers are likely to have had preceding or concurrent low densities of circulating gametocytes. Antibody prevalence in the group classified as gametocyte-negative by microscopy may thus be associated with concurrent low-density gametoctytemia and/or long-lived antibody responses acquired following previous gametocyte exposure. The presently analysed samples thus do not allow any conclusions on a possible role of submicroscopic gametocyte densities in boosting or maintaining antibody responses to gametocyte antigens. Refined studies with longitudinal sampling and gametocyte detection by sensitive qRT-PCR methodologies are needed to formally assess antibody kinetics in relation to gametocyte exposure and determine whether recent markers of exposure to blood stage antigens ^50^ can be complemented by a set of markers for recent or long-term gametocyte exposure.

We described the assembly of a curated gold standard set of gametocyte and asexual proteins and used this new resource to rank the likelihood of all *Pf* proteins and transcripts being specific to the gametocyte stage. Data from 18 publicly available studies were integrated to resolve partially conflicting evidence. The resulting consensus lists can be used for guidance of future investigations as we have shown the value of our predictions by in vitro validation.

## Materials and Methods

### Assembly of a gold standard for gametocyte and asexual proteins to weigh whole proteome/transcriptome data sets

To build a gold standard against which the performance of individual data sets could be assessed, we identified proteins that are known to be expressed in either asexual parasites (mostly blood stage, also including sporozoites and liver stage) or gametocytes. This list was initially informed by literature review (Supplementary table S1) for expression in the respective stages as detected by immunofluorescence assays and/or western blot, supplemented with *P. falciparum* blood stage or transmission blocking vaccine candidates. This initial list was then communicated with experts (including the authors DAB, PA, FS, CJJ, SMK, TWAK, MM, CD, RS and TB) and edited. If additional proteins were suggested for inclusion in the list, published evidence was requested and examined prior to inclusion of the proteinThe final asexual gold standard list contains 46 proteins; the final gametocyte list contains 41 proteins. These gold standard lists (Supplementary Table 1) represent the balance between very strict inclusion criteria and sufficient set size to evaluate the quality of all data sets integrated. We tested for the detection of these proteins or transcripts in the respective samples, using a Bayesian statistics approach that we have successfully applied previously for genes involved in anti-viral immune defense ^31^.

### Data selection and integration

Data sets that measured protein and transcript abundance in *Pf* gametocytes were balanced with data from other life stages and supplemented with studies of the rodent malaria parasite *P. berghei*. One MS study on *P. vivax* was included as it is based on of *ex vivo* blood material as opposed to all other studies that used *in vitro* cultivated parasites. Unique peptide counts were retrieved from plasmoDB (version 28) in which sequenced peptides from the published studies are always mapped to the most recent genome annotation, or supplementary material of the respective studies. Many aspects determine how well a protein is represented in proteomics data that are obtained via MS, like its length or posttranslational modifications. Remapping original MS data to newly annotated genomes improves the quality of the predicted proteins ^19^. We were however not able to retrieve those data from the studies ^13,14,16^ and therefore decided to take those proteins at face value. Notice that also these early studies contribute significantly to our integrated lists. Expression percentiles were retrieved from plasmoDB (version 28) or calculated from raw data in the respective supplementary material. Gametocyte samples were summarized if applicable (using the maximum peptide count/expression percentile of different stages or male and female gametocytes) as were asexual samples, only considering the highest expression in any sample or time point.

For MS and transcriptomics data sets, separate scores for gametocyte-specificity of any *Pf* gene have been calculated. In brief, protein or transcript expression has been categorized from absent to high expression levels as given by number of unique peptides or expression percentiles, respectively. For each of the respective bins, a score was calculated depending on the relative retrieval of gametocyte and asexual gold standard genes. The log ratio of these retrieved genes defined the score for all other genes within the same bin. The final gametocyte score calculates as the prior probability of a gene being gametocyte-specific that is updated using the contributions of the data sets:

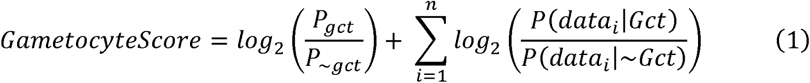

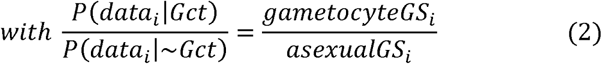

where gametocyteGS and asexualGS are the fractions of retrieved gametocyte and asexual gold standard genes in sample i, respectively. We used a pseudocount of 1 if necessary to prevent division by zero if none of the gold standard genes was retrieved in this specific sample and bin. It was assumed that the likelihood of a gene to be either gametocyte or asexual specific is equally high, thus the (log-transformed) prior equals 0 and the final score depends solely on the integrated data. In the selection of a set of proteins that we assigned to be gametocyte-specific we chose a cutoff score of 5.0 (proteomics-derived). The cutoff score of 5.0 can be interpreted as: a gene has to be 2^5^ = 32 times more likely to be gametocyte-specific than asexual specific. The score of 5.0 was based on the behavior of the gold standard genes. Out of the 41 gametocyte gold standard genes, 37 have a score higher than 5.0, while none of the asexual gold standard genes do.

When applicable, genes from *Pb* and *Pv* were treated as their respective *Pf* orthologs as retrieved from plasmoDB ^54^, to be able to integrate all data sets. When no ortholog is known, the respective non-*Pf* data sets did not contribute to the score of this particular gene. Scores using *Pf* data exclusively were also calculated (Supplementary Table S2 includes all scores and rankings with expression information from all integrated studies).

### Cross-validation of the scoring method

We performed a ten-fold cross-validation to assess the predictive performance of the integrated gametocyte-specificity score (i.e. its ability to discriminate known gametocyte vs. asexual genes). For that, we subsampled both gold standard gene sets ten times (folds), without replacement (i.e. each gene is selected exactly once). Then for each fold we re-weighed and integrated the data sets based on nine-tenth of gold standard genes, and collected the ranks of the one-tenth of genes that were left out in that particular fold. A ROC curve was constructed based on those ranks. Using the same strategy, ROC curves for individual data sets that comprised both gametocyte and asexual samples were constructed for comparison.

### Protein microarray to measure humoral immune responses

A protein microarray that was enriched for gametocyte proteins was produced and probed as described earlier for a study aiming to unravel the immune signature of naturally acquired transmission-reducing immune responses in gametocyte carriers (Stone, Campo *et al*. 2017 accepted manuscript ^36^). As a control group, Gambian asexual parasite carriers without gametocytes detectable by microscopy were included in the probing. For the current study array data from this control group (n=63) and Gambian gametocyte carriers were used (n=164) ^55–60^. All of these 227 individuals were sampled during a period of intense malaria transmission intensity in The Gambia and likely had (multiple) previous malaria infections ^55,61^. For these populations, responses to 30 newly defined highly gametocyte-specific antigens (from 24 genes) were compared between gametocyte carriers and non-carriers (Mann-Whitney U test). Seropositivity for each of the antigens was determined using a mixture model-based cutoff and related to gametocyte carriage using Fisher’s exact test, corrected for multiple testing (Benjamini-Hochberg) of a total of 70 antigens (including 40 antigens from the gametocyte gold standard).

### Transcript abundance in different life stages and strains

The abundance of 15 predicted gametocyte-specific targets was measured in asexual parasites and gametocytes of four different *Pf* strains from *in vitro* culture. The targets were selected from the 100 highest scoring transcripts to account for uncertainties about the absolute scoring of transcriptomics data with a protein-based gold standard. We do not assume a clear hierarchy between these top 100 scoring transcripts and consider any of these genes highly gametocyte-specific. The 15 highest-ranking non-gold standard genes were selected based on a preliminary analysis of the data, and contain genes that are currently not annotated as well as genes with known protein function in gametocytes (PUF1 ^62^ and Ccp4 ^63^). In the final generation of the gametocyte-scores, all validation genes were retained in the top 100 scoring genes. The *Pf* strains used are of West African (NF54, NF166, NF175) and Southeast Asian origin (NF135). All strains were cultured and synchronized as described previously ^20^. Using established standard curves, the same concentrations of parasites were compared for Ct values in qRT-PCR (for primers, see Supplementary Table S7). Extracted nucleic acids were DNase-treated before reverse transcription when introns were absent from the targets. Initial comparison was between mixed asexual blood stage parasites (considering the lowest Ct measured in any strain and replicate) and stage V gametocyte (highest Ct measured per strain and replicate). Promising targets with a high Ct_Asexuals_ - Ct_Gametocytes_ were further examined in serial dilutions of stage V gametocytes and synchronized asexual material of the strain NF54 (10, 20, 30, 40 hours post invasion, resembling early rings, late rings, trophozoites and schizonts, respectively). All qRT-PCR reactions were analyzed in technical triplicates, from biological triplicates (NF54) or duplicates (remaining strains).

RNA samples were used from a clinical malaria trial conducted in Western Kenya ^39^. Samples from days 3 and 7 after treatment were selected to ensure a range of (low-density) gametocyte carriage to test qRT-PCR sensitivity.

### Go term enrichment in top 100 proteins or rank-based enrichment

Current GO term annotation for *Pf* was retrieved from plasmoDB (release 30) and analyzed using the topGO R package ^64^ for the enrichment of terms in the 100 highest scoring proteins versus all *Pf* proteins. Semantic clustering of the significant GO terms in Biological Process ontology was done with the Revigo webtool for *Pf* ^65^. Second, gene set enrichment analysis (GSEA) based on all *Pf* proteins and their ranks and scores was performed using the software available at http://software.broadinstitute.org/gsea/downloads.jsp. Based on the cilium genes reported by the Syscilia consortium ^66^, we assembled a “cilium” GO term of mixed ontology (GO:9999999) for *Pf* with 28 predicted orthologs (Supplementary Table S6).

### Data availability

All data generated or analysed during this study are included in this published article (and its Supplementary Information files).

The gold standard lists (Supplementary Table S1)
Bayesian gametocyte scoring for proteomics and transcriptomics data. Includes *Pf* only-scores and expression values for any gene and individual data set (Supplementary Table S2)
Potential translationally repressed genes with high transcriptomics score (>7) and low proteomics score (<-10, Supplementary Table S3)
Overview over previously reported gametocyte-specificity per study (Supplementary Table S4)
Seroprevalence in The Gambia in gametocyte carriers and non-carriers (Supplementary Table S5)
Function predictions for highly gametocyte-specific proteins with lacking annotation (Supplementary Table S6)
Transcript validation in 15 targets, including primer sequences for qRT-PCR (Supplementary Table S7)
GO term analyses with cilium genes (Supplementary Table S8)

## Acknowledgments

We are thankful to Colin Sutherland (LSHTM London, UK) for commenting on the manuscript and providing serum samples that we used to study antibody responses in Gambian individuals. David Conway and Johannes Dessens (LSHTM London, UK) gave helpful feedback on the initial gold standard lists. We further thank Adam D. Shandling, Jozelyn V. Pablo and Andy A. Teng from Antigen Discovery Inc. (Irvine, CA) for their work on the protein microarray.

TB was supported by the Netherlands Organization for Scientific Research (nwo.nl), through a VIDI fellowship 016.158.306. The Radboud Institute for Health Sciences (rihs.nl), supported LM through grant R-2765. The Virgo consortium (virgo.nl), grant FES0908, supported the work of RvdL and TJPvD. TWAK is supported by the Netherlands Organization for Scientific Research (NWO-VIDI 864.13.009). The funders had no role in study design, data collection and analysis, decision to publish, or preparation of the manuscript.

## Author Contributions Statement

MAH, TB, RvdL and LMK conceptualised the work. LMK, RvdL, TJPvD, MAH and TB analysed the data. LMK, TB, DAB, PA, FS, CJJ, SMK, TWAK, MM and RS assembled the Gold Standard. KL, MvdVB, WG, RSS, LMK and WS conducted and analysed experiments with samples and resources provided by CD and JJC. LMK wrote the first draft of the manuscript, all authors reviewed the manuscript.

